# Spatialproteomics - an interoperable toolbox for analyzing highly multiplexed fluorescence image data

**DOI:** 10.1101/2025.04.29.651202

**Authors:** Matthias Meyer-Bender, Harald Voehringer, Christina Schniederjohann, Sarah Koziel, Erin Chung, Ekaterina Popova, Alexander Brobeil, Lisa-Maria Held, Aamir Munir, Scverse Community, Sascha Dietrich, Peter-Martin Bruch, Wolfgang Huber

**Affiliations:** European Molecular Biology Laboratory (EMBL), Heidelberg, Germany; Molecular Medicine Partnership Unit (MMPU), Heidelberg, Germany; Department of Medicine V, Hematology, Oncology and Rheumatology, University of Heidelberg, Heidelberg, Germany; Department of Hematology and Oncology, University Hospital Düsseldorf, Düsseldorf, Germany; Center for Integrated Oncology Aachen-Bonn-Cologne-Düsseldorf, Düsseldorf, Germany; Spatial & Functional Screening Core Facility, Medical Faculty, Heinrich Heine University, Düsseldorf, Germany; Department of Pathology, University of Heidelberg, Heidelberg, Germany

## Abstract

**Summary:** Highly multiplexed immunofluorescence imaging is a recent method to characterize tissues at single-cell resolution on the protein level, offering low cost, high scalability, and the ability to analyze paraffin-embedded tissue samples. However, the analysis of these data involves a sequence of steps, including segmentation, image processing, marker quantification, cell type classification, and neighborhood analysis, each of which involves a multitude of method and parameter choices that need to be adapted to the data and analytical objective at hand. Moreover, variations in data quality can be high and unpredictable, which necessitates further flexibility and interactivity. While individual components exist, there is an unmet need for a coherent toolbox that offers end-to-end coverage of the workflow, flexibility, and automation.

We present *spatialproteomics*, a Python package that addresses these challenges. Built on top of *xarray* and *dask, spatialproteomics* can process images that are larger than the working memory. It supports synchronization of shared coordinates across data modalities such as images, segmentation masks, and expression matrices, which facilitates easy and safe subsetting and transformation.

We demonstrate *spatialproteomics* on a set of images of reactive lymph nodes or different forms of B cell Non-Hodgkin lymphomas (BNHL) from 132 patients. We showcase an end-to-end analysis from raw images to statistical characterization of cell type composition and spatial distribution across indolent and aggressive lymphomas. Furthermore, we show how *spatialproteomics* can process gigapixel whole slide images. Altogether, we propose spatialproteomics as an easy-to-install, easy-to-learn, comprehensive toolbox for constructing powerful end-to-end image analysis solutions for highly multiplexed immunofluorescence imaging.

**Availability and Implementation:** The source code for *spatialproteomics* is freely available at https://github.com/sagar87/spatialproteomics under the MIT license.

## Introduction

Recent advances in spatial proteomics, such as co-detection by indexing (CODEX) (Black *et al*., 2021), MACSima imaging cyclic staining (MICS) (Kinkhabwala *et al*., 2021), or imaging mass cytometry (IMC) (Giesen *et al*., 2014), enable highly-multiplexed molecular and morphological profiling of tissues at single-cell resolution on a large scale. These techniques generate multi-channel images, where each channel represents the spatial distribution of a distinct protein. Since these assays operate on formalin fixed paraffin embedded (FFPE) tissues, the standard-of-care for archiving diagnostic biopsies, they have found broad adaptation in oncology, where they have been used to characterize the tumor microenvironment (TME) in entities including lymphomas (Roider *et al*., 2024; Phillips *et al*., 2021), colorectal (Li *et al*., 2023), and head and neck cancers (Punovuori *et al*., 2024).

Following Kuswanto et al. (Kuswanto *et al*., 2023), we here consider the following workflow consisting of cell segmentation, image processing, protein quantification and cell phenotyping (**Fig. 1**). This workflow requires the creation of additional data objects, such as segmentation masks and protein expression matrices. While these, as well as the primary imaging data itself, could be represented simply as an unstructured collection of multidimensional arrays, many of them share common dimensions (**Sup. Fig. 1**). For example, the multiplexed imaging data (channels, x, y) share spatial dimensions with their corresponding segmentation masks (x, y). Similarly, the instance labels in a segmentation mask can be viewed as a dimension that aligns with the expression matrix (cells, channels) quantifying protein expression for each cell, or with a morphology matrix (cells, shape features) of shape and texture descriptors for each cell. Therefore, a desirable data structure for multiplexed imaging data should keep such shared dimensions consistent, ensuring that changes to one component automatically update the others to maintain consistency. For example, when subsetting a region of an image, the segmentation mask should be subset accordingly, and the expression matrix should only retain cells within this region. Similarly, it should be possible for users to select specific cell types and forward the resulting subset object to downstream methods for the computation of spatial statistics.

**Figure 1:**
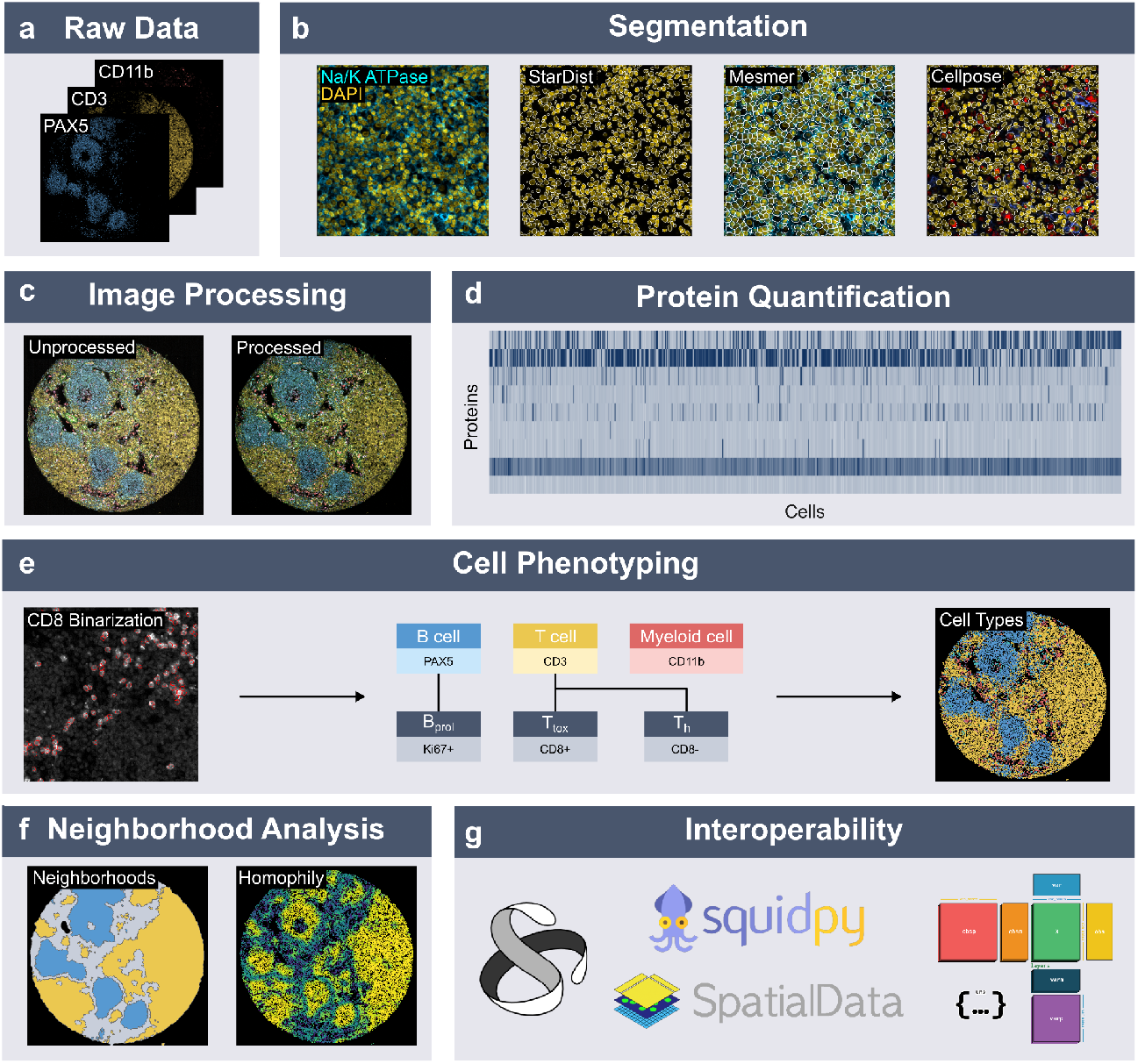
An exemplary workflow orchestrated with *spatialproteomics*. **a-b**, After raw images are obtained, segmentation is performed either on the nuclear or whole cell level, or on multiple markers independently. **c**, Image processing tools such as thresholding can boost the signal-to-noise ratio of the image. **d**, Fluorescence intensities are quantified for each segmented cell. **e**, The resulting expression matrix serves as input for cell type prediction, both of major cell types and subtypes. **f**, These cell type predictions can then be used to perform neighborhood analysis or find areas of tissue heterogeneity. **g**, Export into data formats like *anndata* and *spatialdata* ensures interoperability with the *scverse* ecosystem.

We introduce *spatialproteomics*, a Python package that leverages *xarray (Hoyer and Hamman, 2017)* to provide a consistent representation of multiplexed imaging data and associated data types across shared dimensions. The package offers a unified API for image processing, segmentation, marker quantification, cell type classification, neighborhood analysis and visualization. Functions in *spatialproteomics* are endomorphisms on the *xarray* class, which facilitates the “piping” of the data through a series of such operations by function composition (Bird, 1998). Furthermore, *spatialproteomics* can process high-dimensional larger-than-memory imaging datasets in parallel via *dask (Rocklin, 2015)*.

S*cverse* is an open-source ecosystem of interoperable data structures and tools for analyzing single-cell omics data (Virshup *et al*., 2023). *Spatialproteomics* integrates seamlessly into the *scverse* framework by supporting import and export to both the *anndata* and *spatialdata* formats (Virshup *et al*., 2021; Marconato *et al*., 2024). Additionally, by working directly with *spatialdata* objects, *spatialproteomics* can be incorporated into established workflows within the *scverse* ecosystem. As one of the first domain-specific packages to integrate with both *anndata* and *spatialdata*, it contributes to expanding the ecosystem toward spatial proteomics and facilitates adoption through standardized interfaces and community-aligned design. The package is available at https://github.com/sagar87/spatialproteomics.

## Results

### Spatialproteomics streamlines the processing of highly multiplexed fluorescence images

We considered a dataset of lymph node biopsies from patients diagnosed with B cell Non-Hodgkin lymphoma (BNHL). Tissue microarrays (TMAs) and whole slides were imaged using a set of 56 antibodies selected for specific binding to proteins important in B, T and myeloid cell biology (Roider *et al*., 2024). We obtained 250 TMA tiles with a size of 9 megapixels (3000×3000 pixels) each, and five whole slide images (WSIs) with sizes roughly between 1 and 2 gigapixels (**Sup. Table 1**). We designed *spatialproteomics* as a toolbox to rapidly put together and adapt end-to-end analysis workflows for large scale datasets from different tissues and using different antibody panels and methods.

To load the data into Python, *spatialproteomics* requires tiff files and a mapping from each channel to its corresponding protein, as is the standard output of many of the above listed techniques. For visualization, users can select which channels to show and assign custom colors to each channel (**Fig. 1a**). After data import, we performed quality control by visually inspecting the images. To facilitate interactive work and conserve memory, users can subset or downsample images.

Functions in *spatialproteomics* follow an endomorphic design pattern, allowing commands to be chained seamlessly and eliminating the need for intermediate variables or tedious data structure conversions. This improves code readability, reduces memory usage, and ensures consistent synchronization of related data. Since all functions in the package adhere to this pattern, complex analysis pipelines can be constructed in just a few lines of code.

The assessment of data quality is followed by cell segmentation (**Fig. 1b**). *Spatialproteomics* offers wrappers for *stardist* (Schmidt *et al*., 2018), *mesmer* (Greenwald *et al*., 2022), and *cellpose* (Stringer *et al*., 2021), enabling rapid experimentation with different segmentation methods. All of these employ neural networks that were pre-trained on thousands of images, and provide segmentation of nuclei and optionally also of whole cells. Users can also load their own segmentation mask directly into the *spatialproteomics* object, making the workflow adaptable to new methods.

External tools can have incompatible dependencies, sometimes rendering it impossible to install all of them within the same virtual environment. While *spatialproteomics* offers the option to install all external tools during the installation process, users can also choose to install only a subset to minimize potential version conflicts.

In some cases, segmentation on a nuclear or a universal membrane marker does not provide satisfactory results, and more elaborate approaches are required. For example, segmentation of macrophages or dendritic cells can be hampered by their large size beyond the imaged tissue’s thickness and irregular shape. To counteract this problem, we can perform independent segmentation on several markers, e.g., DAPI (nuclei), CD68 (macrophages) and CD11c (dendritic cells). The resulting masks can then be merged to obtain an improved segmentation.

Antibody-based imaging can suffer from high background (low signal-to-noise ratio) and high variability of binding efficiency and specificity across samples or even within the same image. To address such issues, users can apply predefined or custom-written image processing steps, such as thresholding, filtering, and morphological operations (Russ, 2006) (**Fig. 1c**).

Following segmentation and optional image processing, an expression matrix is computed by summarizing intensities over each cell mask (**Fig. 1d**). *Spatialproteomics* allows different summarization functions, including average, sum, median, percentage of non-zero pixels; users can also provide their own summary functions. To facilitate downstream analysis and comparisons across markers and samples, transformations such as min-max normalization, z-score or arcsinh-transform can be applied to the resulting expression matrix (Hickey *et al*., 2021).

Many downstream analysis approaches benefit from annotating the observed cells with cell type labels. Such cell type classification is, however, a bespoke task for which there are no general or “out-of-the-box” solutions. It is contingent both on the scientific goals of the study, which can direct the granularity of the cell type classification, and on technological limitations. *Spatialproteomics* facilitates the creation of such custom-made cell type annotation workflows. We advocate a decision tree approach (**Fig. 1e**). This requires the user to define a set of mutually exclusive marker proteins for each of the cell types or states they want to distinguish and annotate. The approach is analogous to gating methods used in multi-dimensional flow cytometry (Verschoor *et al*., 2015). After thresholding the image for those markers to remove unspecific background signal, we predict cell types based on identifying the cell type whose corresponding marker expression is largest.

This has two advantages:

1. A particularly challenging technical artefact is so-called *spillover*, in which protein expression signals from a neighbouring cell are counted towards the expression profile of the currently considered cell. A prominent reason for spillover is segmentation errors, or more fundamentally, uncertainties in the assignment of particular pixels to one cell or another.
2. With unsupervised clustering methods, the identified clusters need to be annotated post hoc based on marker expression profiles. Techniques that incorporate prior knowledge about existing cell types and their corresponding markers a priori circumvent this step, so that such associations only need to be defined once.

The second step of cell phenotyping consists of partitioning the major cell types into subtypes. Some proteins, such as the proliferation marker Ki-67, can be expressed by multiple different cell types. To enable the partitioning of major cell types into a more fine-grained classification scheme, such markers can be binarized, resulting in each cell being “positive” or “negative” for a given marker.

*Spatialproteomics* allows users to define a hierarchical cell type gating tree, which partitions the previously predicted major cell types into more fine-grained subtypes. This scheme also enables more complex marker combinations. Next to single marker positivity or negativity, a cell subtype can also be defined by a combination of both positive and negative expression, or by a set of alternative markers (e. g. a cytotoxic T cell could be predicted to be exhausted if it expresses either PD1 or TIM3).

Next to our hierarchical gating method, we also implement a wrapper around *assignment of single-cell proteomics* (astir) (Geuenich *et al*., 2021). This method offers a probabilistic approach to cell type prediction and also provides uncertainty estimates about its predictions.

A commonly used approach to detect changes in tissue structure when comparing conditions is the definition of cellular neighborhoods (Schürch *et al*., 2020; Kim *et al*., 2022; Varrone *et al*., 2024; Kuswanto *et al*., 2023; Nirmal and Sorger, 2024) (**Fig. 1f**). In this approach, a neighborhood profile gets computed for each cell, which consists of the relative cell type abundances surrounding the center cell. These neighborhoods can be defined via Delaunay triangulation, the k nearest neighbors or a circle with a fixed radius centered at each cell. Having obtained one neighborhood profile per cell, unsupervised clustering such as k-means clustering can reveal condition-specific tissue composition.

*Spatialproteomics* allows users to investigate neighborhoods in more detail. Since the neighborhood of a cell can be modeled as a network, where each cell is connected to all cells within its vicinity, a variety of features can be computed for each cell. Measures such as homophily (the fraction of cells in the neighborhood with the same cell type as the center cell) and Shannon’s diversity index can provide information about how homogeneous the tissue is around any cell. The definition of cellular neighborhoods also enables users to subset cells, for example only selecting cells belonging to a tumor region or cells with a homogeneous microenvironment. Further investigation of the neighborhood composition and heterogeneity allows for a more detailed description of the tissue.

Putting everything together, *spatialproteomics* provides static plotting methods to not only visualize marker intensities, but also segmentation masks, cell type labels and neighborhoods. On disk, objects are stored as *zarr* files, so that users can load images, segmentation masks, quantifications, cell type predictions and other possibly interesting data without needing to store the whole object in memory. Additionally, we implemented export functions to *AnnData* (Virshup *et al*., 2021) and *spatialdata (Marconato et al*., *2024)* (**Fig. 1g**), which can in turn be used with spatial analysis tools such as *squidpy* (*Palla et al*., 2022), SCIMAP (*Nirmal and Sorger, 2024)*, and *Multiomics and Ecological Spatial Analysis* (MESA) (Ding *et al*., 2025). This interoperability also enables interactive analysis using tools such as *napari* (Sofroniew *et al*., 2024).

### Spatialproteomics enables the large scale analysis of a diverse dataset of B cell Non-Hodgkin lymphomas

To demonstrate the capabilities of *spatialproteomics*, we performed spatial profiling on 250 samples of 132 patients with B cell Non-Hodgkin lymphomas (BNHL) or non-neoplastic controls. Lymph node biopsies were extracted from the patients, and tissue microarrays (TMA) were created using 1mm diameter cores and a median of two replicates per patient. The samples were grouped into three distinct categories based on their condition: reactive lymph nodes (LN), indolent lymphomas with retained tissue structure (marginal zone lymphoma (MZL), follicular lymphoma (FL)), and aggressive lymphomas with diffuse structure (diffuse large B cell lymphoma (DLBCL), primary mediastinal large B cell lymphoma (PMBCL), Burkitt’s lymphoma, B cell lymphoblastic lymphoma (BLBL)). We imaged 56 markers with the purpose of inferring cell types and states from their expression patterns (**Sup. Table 2**). In total, more than 3.5 million cells were segmented, with the average number of cells detected per core being 14,273.

BNHL consist of malignant B cells along with various other cell types within the tumor microenvironment (TME). Although the composition of the TME is heterogeneous across disease subtypes, several major cell types, such as T cells, macrophages, and endothelial cells, are commonly present (Scott and Gascoyne, 2014). These can be subdivided further according to the expression of additional lineage-specific or functional markers. Previous studies showed that the frequencies of specific T cell subpopulations vary across BNHL subtypes - partly due to differences in clonal expansion - and can be associated with survival (Roider *et al*., 2024). To explore this, we constructed a hierarchical gating tree that focused specifically on the classification of T cell subsets (**Fig. 2a**).

**Figure 2:**
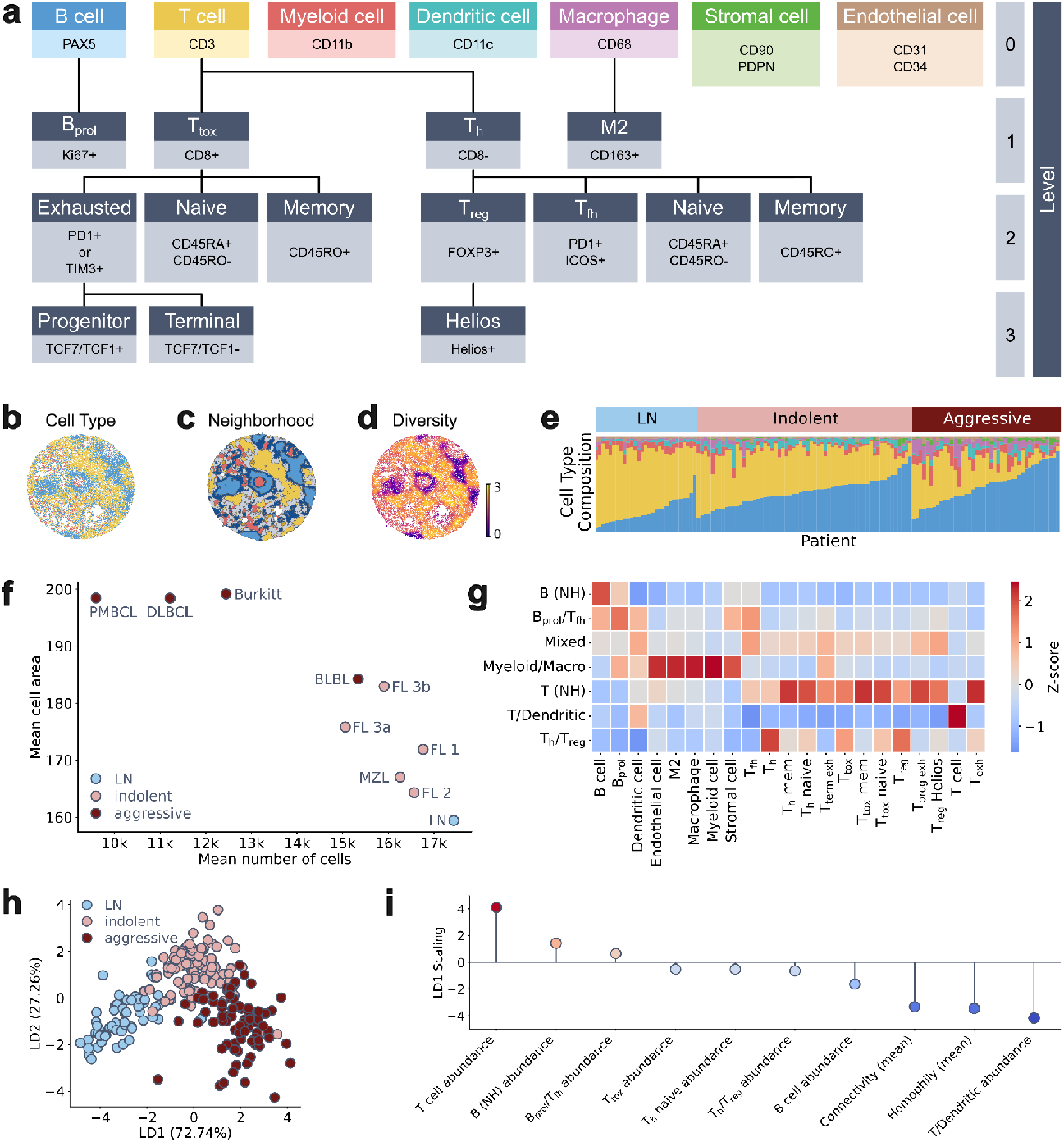
Analysis of a diverse selection of B cell Non-Hodgkin lymphomas. **a**, A hierarchical gating tree used for cell type and subtype prediction. **b-d**, Cell types, neighborhoods, and Shannon’s diversity index of a single lymph node core. Only coarse cell types are shown in the visualization. Cell type colors correspond to the ones shown in panel a. **e**, Overview of patients and their cell type compositions, cell type colors as specified in panel a. **f**, More aggressive lymphomas tend to have larger cells than lymph nodes. **g**, Clustered neighborhoods (NH) capture distinct cell type compositions. **h**, Linear discriminant analysis (LDA) of samples based on cell type abundances, neighborhood abundances, and morphological information. **i**, LD1 scalings of the individual features in the space spanned by the class centroids from the LDA. Only features with an absolute scaling above 0.5 are shown.

We predicted the cell type of each individual cell according to this gating tree (**Fig. 2b**). A comparison of cell type composition showed that lymphoma samples had a significantly higher content of B cells (**Fig. 2e, Sup. Fig. 2**), which is expected due to accumulation of malignant B cells. Furthermore, we noted an increase in macrophages and stromal cells pointing towards increased vascularization of the tumor samples (**Sup. Fig. 2**). A principal component analysis based on relative cell type abundances corroborated this finding, showing that the main axis of variation was driven by B, T, and stromal cells (**Sup. Fig. 3**).

Next to the cell type composition, we investigated cell areas in the different conditions. Aggressive lymphomas showed larger cells, while the average cell area in indolent lymphomas was closer to the reactive lymph nodes (**Fig. 2f**), in line with the blast-like morphology of aggressive lymphoma (Alaggio *et al*., 2022). As a consequence, cores from aggressive lymphoma contained fewer cells in the same tissue area.

Beyond looking at individual cell types, we collected the cell type composition within a radius of 25 µm around each cell and clustered them across samples to obtain shared neighborhood profiles (**Fig. 2c**,**g**). We identified seven distinct neighborhoods, including a B cell rich neighborhood (B (NH)) and one enriched in proliferating B cells (B_prol_/T_fh_). Another neighborhood consisted mainly of myeloid cells and macrophages (Myeloid/Macro), and one showed enrichment in T helper and regulatory T cells (T_h_/T_reg_). The “T/Dendritic” neighborhood predominantly captured T cells that could not be further classified due to limited CD8 signal. While this reflects a limitation of the hierarchical cell type classification in low signal-to-noise settings, we observed this neighborhood in only a few samples (**Sup. Fig. 6**).

Given cell type abundances, neighborhood abundances, as well as morphological information such as the mean homophily of every patient, we were interested to see which of these metrics could be used to differentiate between lymph nodes, indolent and aggressive lymphomas. We performed a linear discriminant analysis on the three groups, which showed a clear trajectory from reactive lymph nodes over indolent to aggressive lymphomas (**Fig. 2h**). To see which features were most important for the classification, we considered the scaling of each feature in the space spanned by the class centroids (**Fig. 2i**). Aside from cell type abundances, one major driving factor was homophily, which is in line with observed diffuse architecture in most aggressive lymphomas, showing that there is merit to obtaining features in addition to cell type abundances.

### Spatial statistics reveal clustering of cell types and cell-to-cell interactions

While cellular neighborhoods provide one perspective into the spatial distribution of cell types, there are many more spatial statistics methods to further investigate and characterize tissue architectures, cell-to-cell interactions, and higher-order patterns. We set out with the goal of identifying which cell types occurred more frequently in clusters and which ones were randomly spatially distributed. To achieve this, we computed Ripley’s K function with a maximum radius of 150 µm and looked at the area between the observed curve and the theoretical distribution that would emerge if the spatial point pattern was generated by a homogeneous Poisson process. **Fig. 3a** shows that B cells, proliferating B cells, and T follicular helper cells occur in clusters in reactive lymph nodes, corresponding to germinal centers. As tissue structure becomes more dispersed in aggressive lymphomas, the spatial clustering of these cell types becomes less prevalent.

**Figure 3:**
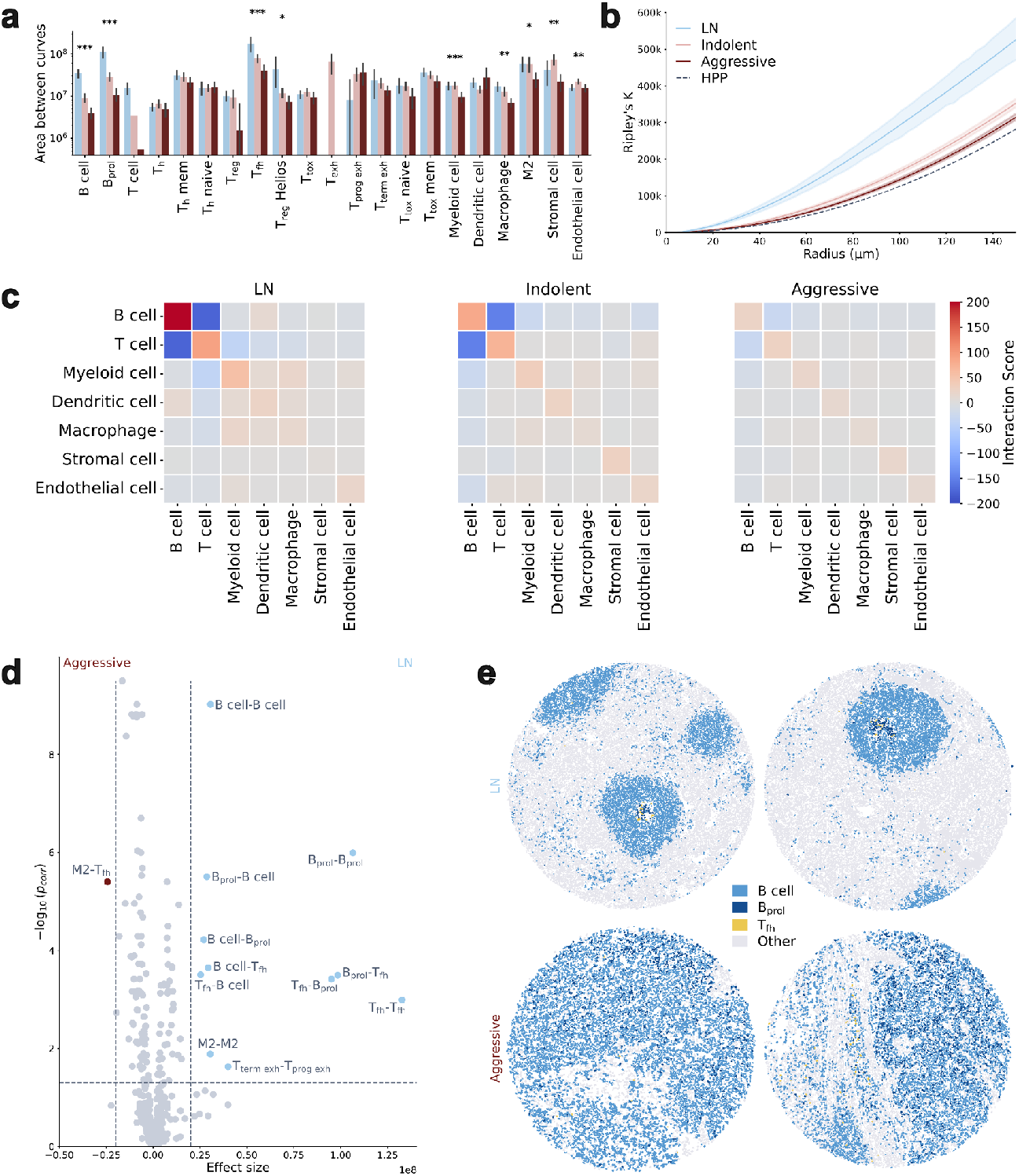
Spatial analysis of the BNHL cohort. **a**, Area between theoretical and observed Ripley’s k. Significance determined via ANOVA. Some cell types (B cells, B_prol_, T_fh_, …) cluster more in lymph nodes and lose structure in lymphomas. **b**, Ripley’s K for B cells, resolved by entity class. The black line shows the expected distribution if the point distribution followed a homogeneous Poisson process (HPP). **c**, Mean interaction z-scores from *squidpy* on the level 0 of the gating tree. Interactions diminish in more aggressive lymphomas. **d**, Ripley’s K cross on the fine-grained cell types. Differences between LN and aggressive, p-values determined with an independent t-test. **e**, Representative cores for LN and aggressive lymphomas show that spatial organization gets lost.

Next to cells of the same cell type clustering together, we wondered if there are certain combinations of cell types which co-occur frequently. To investigate this, we considered a distance of less than 70 µm between cell centroids and computed the neighborhood enrichment score using *squidpy* (*Palla et al., 2022*). We applied this to the level 0 cell types, which showed that the structure becomes more diffuse with aggressiveness of the lymphomas (**Fig. 3c**).

To consider the fine-grained cell type annotations, we computed the areas between the observed and theoretical Ripley’s Cross-K function for each pair of cell types, and compared them between lymph nodes and aggressive lymphomas (**Fig. 3d**). As before, the majority of significant differences were interactions that were present in lymph nodes and disappeared in the lymphoma samples. While most of these interactions were between cells of the same cell types, some interactions reminiscent of germinal centers were also observed, such as between proliferating B cells and T follicular helper cells. On the flipside, an interaction between M2 macrophages and T follicular helper cells seemed to be enriched in aggressive lymphomas.

### Validation using five DLBCL whole-slide images

TMA cores only show a small part of the underlying tissue, and it is typically assumed that they show a representative excerpt of the whole tissue. To see how well our TMAs recapitulate the overall tumor architecture, we performed whole slide imaging on five samples from patients with diffuse large B cell lymphoma (DLBCL), for which we had two matched TMA cores each. After segmentation with cellpose and the removal of cells which were not part of the main tissue of interest, we obtained a dataset of over 6.6M cells, which is almost twice as much as all of the previous TMAs had together.

Despite *Akoya*’s built-in illumination correction using the *BaSiC* algorithm (Peng *et al*., 2017), we still observed differences in marker intensities across tissues, hampering the application of our thresholding approach. To adjust for these, we leveraged *spatialproteomics’* ability to apply arbitrary image processing methods to the multi-channel images. For each channel, we constructed a second image that was a Gaussian blur of the original image, with a kernel size of 51 pixels, and subtracted it from the original one (**Fig. 4a**). This reduced the image-wide gradients, making it possible to apply thresholding again and use the processed images to predict cell types as previously described (**Fig. 4b**).

**Figure 4:**
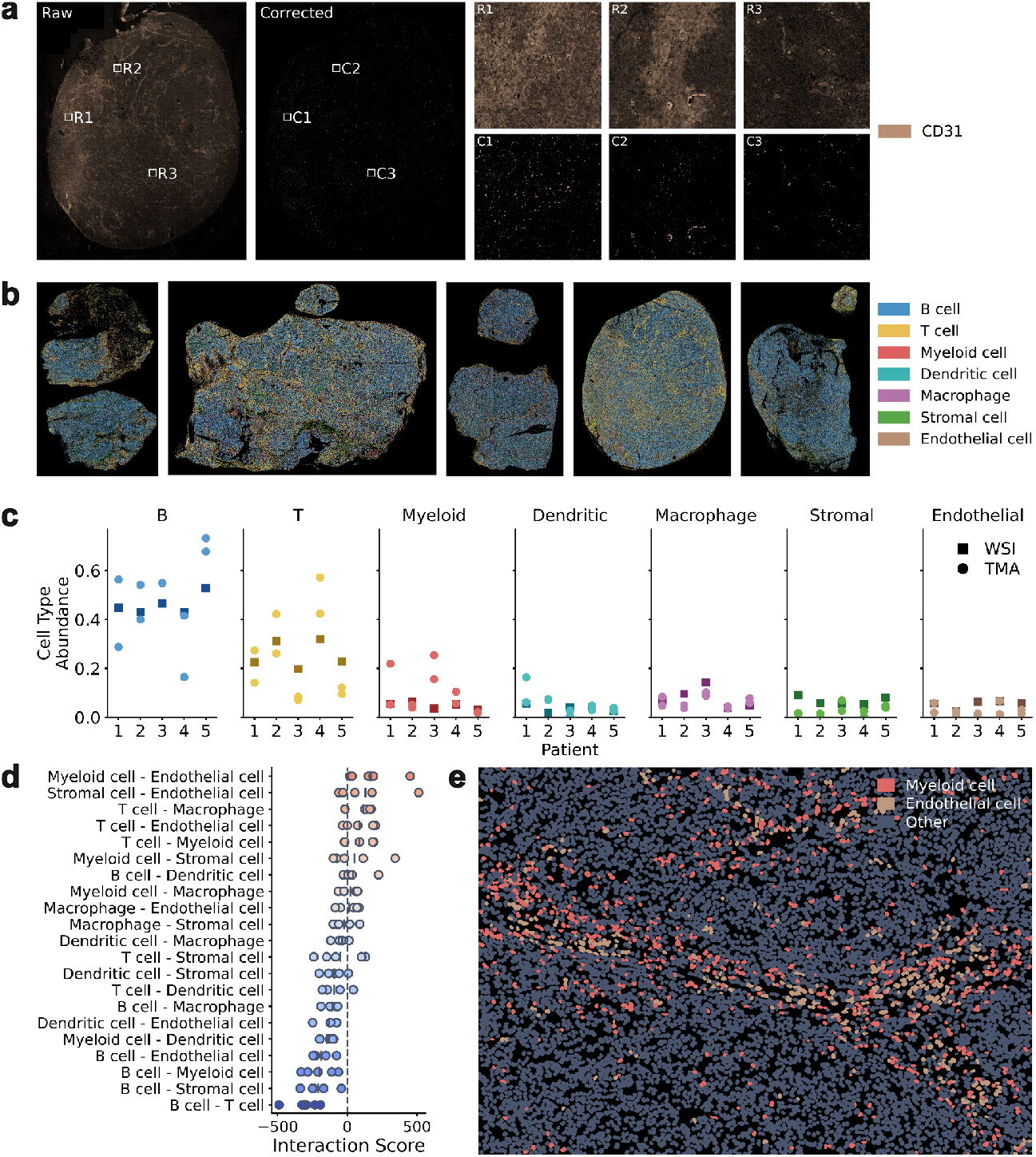
Analysis of five DLBCL whole slide images. **a**, Spatialproteomics enables the application of arbitrary image processing algorithms to multi-channel images, allowing us to correct for differences in marker intensities across whole slide images (WSIs). **b**, Predicted cell types. **c**, Comparison of cell type abundances between whole slide images and corresponding TMA cores. **d**, Interaction scores from squidpy’s neighborhood enrichment function, excluding self-interactions. **e**, Example of an interaction between myeloid cells and endothelial cells.

Next, we were interested in comparing cell type abundances between the WSIs and their corresponding TMA samples (**Fig. 4c**). While the general trend in cell type composition with many B and T cells held true in the whole slide dataset, there were some discrepancies to the TMA results. For example, the WSI for patient 2 contained some connective tissue, which was mostly classified as myeloid cells and therefore skewed the ratio of cell types.

To make use of the spatial context, we again applied the neighborhood enrichment method from *squidpy* (**Fig. 4d**). For clarity, we only considered interactions between different cell types, since self-interactions were frequently most enriched. In accordance with our TMA results, B cells and T cells had a negative interaction score. We also found several enriched interactions such as Myeloid-Endothelial, Stromal-Endothelial, and T-Macrophage (**Fig. 4e**). While these interactions were computed without taking into account a negative control, they do show that there are interactions between specific cell types in DLBCL, providing opportunities for future research.

## Discussion

Highly multiplexed immunofluorescence imaging enables molecular profiling of cells in their native spatial organization in tissues. Analyzing such data requires multiple processing steps that generate additional, derived layers of data – such as segmentation masks, marker intensities, and cellular annotations – all of which share some common dimensions, but may also introduce new ones. Each step can build upon a rich diversity of established methods and existing software implementations. A main task of the analyst is to construct end-to-end workflows that seamlessly chain the steps together, adapting and optimally tuning each of them. Here we present *spatialproteomics*, an open-source Python package that provides a comprehensive toolkit based on a unified data structure that maintains synchronization across dimensions throughout.

We have drawn inspiration and used experience from several existing tools. In the R ecosystem, *cytomapper* and *imcRtools* (Windhager *et al*., 2023) build upon the *SpatialExperiment* data structure (Righelli *et al*., 2022) for spatial single-cell data. In Python, *Spatial Omics Pipeline and Analysis* (Sopa) (Blampey *et al*., 2024) and *Structured Spatial Analysis for Codex Exploration* (SPACEc) (Tan *et al*., 2024) offer similar functionality for processing and analyzing highly multiplexed fluorescence images. Additionally, pipelines such as *Multiple Choice MICROscopy* (MCMICRO) (Schapiro *et al*., 2022) and *Single-cell Identification from MultiPLexed Images* (SIMPLI) (Bortolomeazzi *et al*., 2022) provide Nextflow-based solutions.

The package presented here, *spatialproteomics*, wraps widely used implementations of methods for segmentation, marker quantification, cell type prediction, neighborhood analysis, and visualization. By unifying these methods under a consistent application programming interface (API), *spatialproteomics* enables researchers to experiment with different method choices and parameters when they adapt a workflow to their data and scientific questions of interest. A standardized syntax further facilitates exploration of diverse analytical approaches. Importantly, *spatialproteomics* automatically manages shared dimensions between data layers. This relieves the user of tedious book-keeping and eliminates a frequent source of – sometimes hard-to-find – errors. The package is designed for easy installation, and integrates seamlessly with the *scverse* ecosystem.

We showed an application of *spatialproteomics* to a dataset of BNHL biopsies and found that some of the derived features could effectively differentiate between non-malignant lymph nodes, indolent lymphoma, and aggressive lymphoma. The results recapitulated the loss of germinal centers and overall tissue structure in diseased lymph nodes. We validated some of the results on whole slide images, further demonstrating the scalability of *spatialproteomics* for handling large datasets.

S*patialproteomics* provides access to a range of spatial processing and analysis methods. It seamlessly integrates into the existing single-cell data analysis infrastructure of the *scverse* ecosystem. By lowering technical barriers and aligning with familiar idioms from single-cell analytics, we hope that our package will help spread and increase the utility of spatial proteomics in biomedical and basic biology research.

## Methods

### Sample Preparation

Microscopy slides with 5 µm sections of the BNHL TMAs and DLBCL whole-slide images were prepared by the National Center for Tumor Diseases (NCT) tissue bank in Heidelberg (ethics vote S-686/2018). Multiplex immunofluorescence was performed as previously described (Schniederjohann *et al*., 2025). In brief, the tissue samples were deparaffinized and re-hydrated. Heat-induced antigen retrieval was performed for 20 min at 155-160°C using Tris-EDTA buffer at pH 9. To reduce autofluorescence, tissue samples were bleached two times for 45 min using hydrogen peroxide (H2O2). Tissue samples were incubated with a cocktail of 56 DNA oligonucleotide-conjugated antibodies overnight at 4°C (**Sup. Table 2**). For antibody conjugation, DNA oligonucleotide sequences previously published were used (Schürch *et al*., 2020). The next day, a three-step fixation followed including incubation with 1.6 % paraformaldehyde (PFA), 100 % methanol and bissulfosuccinimidyl suberate (BS3).

Tissue samples were imaged using the Phenocycler Fusion System (Akoya Biosciences) according to the manufacturer’s instructions. For cyclic imaging, Atto550 and Atto647N labelled complementary DNA-barcodes were used. Acquired raw files were processed into a single qpTIFF file by the Fusion software (version 1.0.7 for TMAs, version 2.2.0 for WSI). Details on the raw data processing are available in the manufacturer’s technical documentation (https://help.codex.bio/codex/processor/technical-notes/). Briefly, for each cycle, the nuclear stain pattern was compared to the reference cycle - by default, the DAPI signal from the second cycle. The calculated offsets were used to align all cycles through rigid translation.

Background subtraction was performed using “blank” cycles at the beginning and end of each experiment. Deconvolution was carried out using the *Microvolution* package, while shading correction was applied with *BaSiC*. Image registration of adjacent tiles was performed using the *Microscopy Image Stitching Tool* (MIST) library.

### TMA Cropping

A grid defining the locations of the cores within the tissue microarrays was manually constructed in QuPath using the built-in TMA de-arrayer. The coordinates were exported to a csv, and crops of size 3,000 by 3,000 pixels were extracted around the centroid of each crop.

### Segmentation

For the TMAs, cells were segmented on the DAPI channel using the *cyto3* model from *cellpose 3*.*0*.*7*, and filtered to exclude cells with an area below 50 pixels or above 300 pixels. Subsequently, masks were expanded into each direction by two pixels.

### Quality Control

We manually excluded 28 samples from the original dataset due to artefacts such as tissue dissociation, marker spillage, or other contamination of the image. We also only kept samples for which clinical annotations were available. For samples where only part of it was corrupted by an artefact, we manually drew in masks and removed all cells within those masks from downstream analysis.

On the cell level, cells which were too isolated from other cells were removed. We achieved this by running a binary dilation of 25 pixels on the segmentation masks, finding connected components and removing components which contained less than five cells.

### Image Processing and Cell Phenotyping

For the classification of major cell types, we first thresholded the different markers by quantiles, depending on the specificity and overall signal intensity of each marker. We started with the following default values: {PAX5: 0.8, CD3: 0.7, CD11b: 0.8, CD11c: 0.9, CD68: 0.9, CD31: 0.95, CD34: 0.95, CD90: 0.95, Podoplanin: 0.95}. We then manually cross-checked the images with the cell type predictions, and adjusted the thresholds where we deemed necessary (**Sup. Table 3**). The thresholding itself consisted of computing the corresponding quantile value, subtracting it from the original image, and then clipping the result to 0 to remove negative values.

After thresholding on the pixel level, we aggregated the marker intensities by taking the mean over each segmentation mask. We then applied an arcsinh transform with a cofactor of 5 to the resulting expression matrix. Finally, we assigned the cell type whose corresponding marker had the largest expression. The correspondence between cell types and markers is shown in **Fig. 2a** (level 0).

For the functional markers, we once again thresholded the markers manually, and then computed the percentage of positive pixels which remained within each segmentation mask after thresholding. Using a second threshold t, cells with t% or more positive pixels were assigned as “positive”, while all others were considered “negative” (**Sup. Table 3**).

In some samples, certain markers were too unspecific to enable reliable thresholding. In such cases, we stopped classification at the corresponding node of the gating tree (**Fig. 2a**). For example, when we were not able to binarize CD8, we simply called all T cells as “T cells”, and did not traverse the gating tree further. The advantage of this was that we could retain samples even if an individual marker did not exhibit specific staining characteristics.

### Neighborhood Analysis

To define neighborhoods, we considered cells whose centroids were within 25 µm from one another. After quantifying the cell type abundances within the 25 µm radius around each cell, we applied k-means clustering with k = 7 to obtain neighborhood labels. The numerical labels were then manually substituted by more informative names based on the composition of each cluster (**Fig. 2g**).

### Morphological Profiling of Niches

We implemented various metrics to quantify the heterogeneity of the environment around each cell. These metrics include:

1. Degree: the number of cells connected to each cell
2. Homophily: the fraction of neighboring cells which have the same cell type as the center cell
3. Inter-Label Connectivity: the fraction of neighboring cells which have a different cell type than the center cell
4. Shannon’s diversity index: 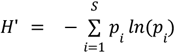, where *p*_*i*_ are the cell type proportions.

To compare these between samples, we computed the mean across all cells within a sample.

### Spatial Statistics

To investigate the clustering behaviour of individual cell types, we used the *spatstat* package in R to compute Ripley’s K function in each sample on each cell type. We then limited the distance to 150 µm and computed the area between the observed distribution and the theoretical distribution that would emerge if the point pattern was generated by a homogeneous Poisson process.

To show the average lines in a single plot (**Fig. 3b**), we interpolated the data along 100 points between the distances of 0 and 150 µm using one-dimensional linear interpolation.

For *squidpy*, we used a radius-based definition of neighborhoods, defining two cells to be interacting if their centroids were less than 70 µm apart.

To further investigate cell-cell interactions, we also computed Ripley’s K cross using the *spatstat* package. We then computed the area between the observed and theoretical curves between 0 and 150 µm, and performed an independent T-test to evaluate the differences between lymph nodes and aggressive lymphomas (**Fig. 3d**). We used Benjamini-Hochberg correction with a family-wise error rate of 0.05 to adjust the p-values. Interactions were deemed significant if their FDR-corrected p value was below 0.05, and the effect size was above 20M (**Fig. 3d**).

### Analysis of Whole Slide Images

We segmented the whole slide images using the *cyto3* model from *cellpose* 3.1.0 and removed cells whose area was outside of the range between 50 and 600 pixels. In addition, we drew regions of interest around the main tissue to remove outlying or irrelevant cells.

To avoid marker gradients skewing the cell type prediction, we applied a background correction by creating a copy of the original image, applying a Gaussian blur with a kernel size of 51 pixels, and subtracting the result from the original image. Cell type prediction was performed as before by setting a threshold for each marker and assigning the cell type with the highest expression per cell (**Sup. Table 3**).

## Code Availability

*Spatialproteomics* can be installed via pip, and the code is publicly available on GitHub at https://github.com/sagar87/spatialproteomics under the MIT license. Documentation can be found under https://sagar87.github.io/spatialproteomics.

## Acknowledgements

We thank the Tissue Bank of the National Center for Tumor Diseases and Institute of Pathology at the University Hospital Heidelberg, who provided biomaterials for the multiparametric immunofluorescence analyses. We thank all members of the Huber group and the Dietrich lab for their support and discussions. We also thank the *scverse* community and all collaborators who provided early feedback on the *spatialprotemics* package.

## Supplementary Information

**Supplementary Figure 1:**
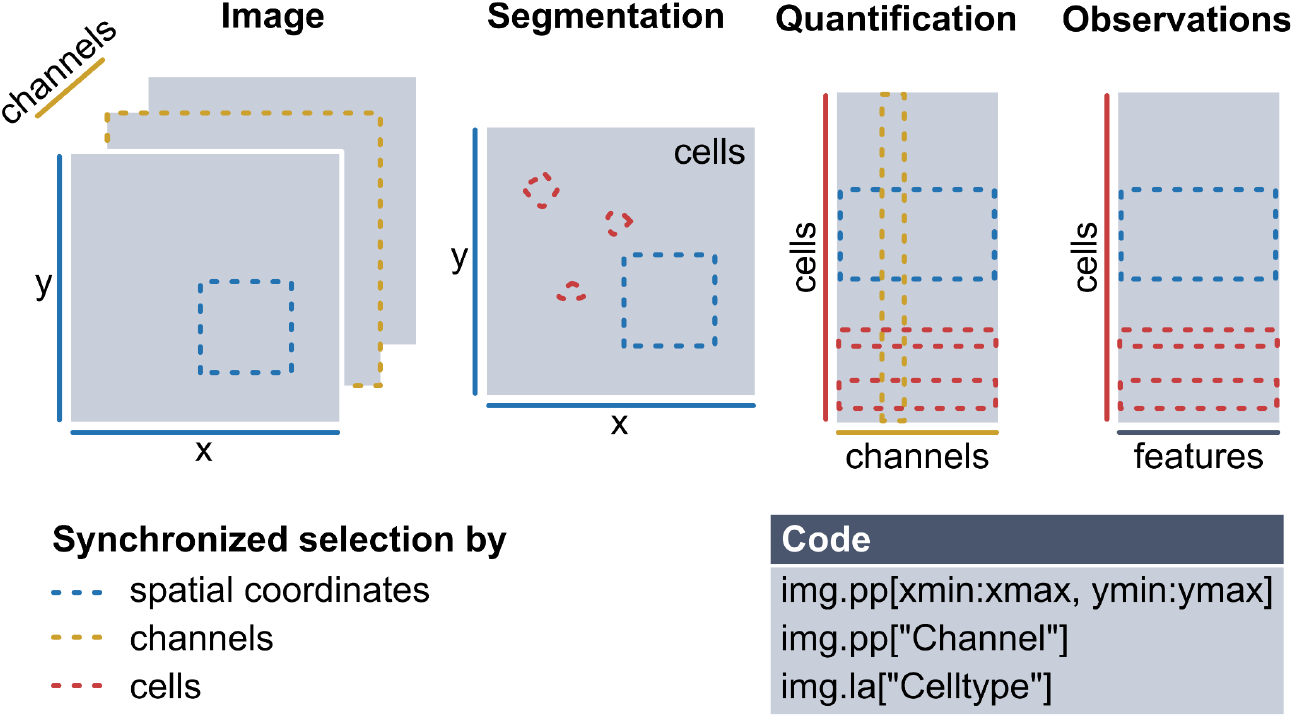
S*patialproteomics* ensures synchronization of data with shared dimensions. Users can subselect data based on spatial coordinates, channels, cell types, and more.

**Supplementary Figure 2:**
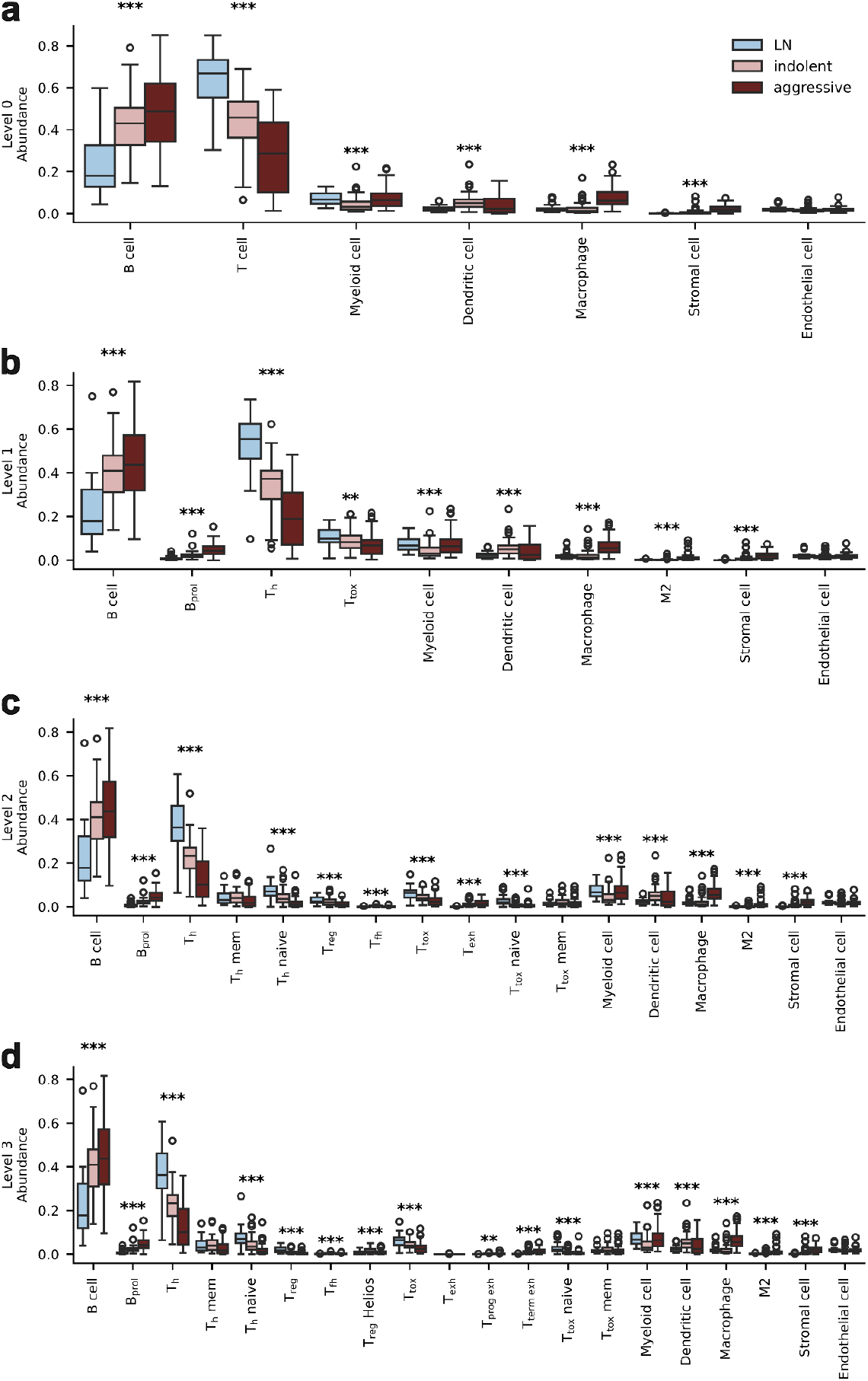
Cell type abundances quantified at different granularities of the gating tree. Statistical significance was assessed using ANOVA. Significance levels are indicated as follows: p < 0.05 (*), p < 0.01 (**), and p < 0.001 (***).

**Supplementary Figure 3:**
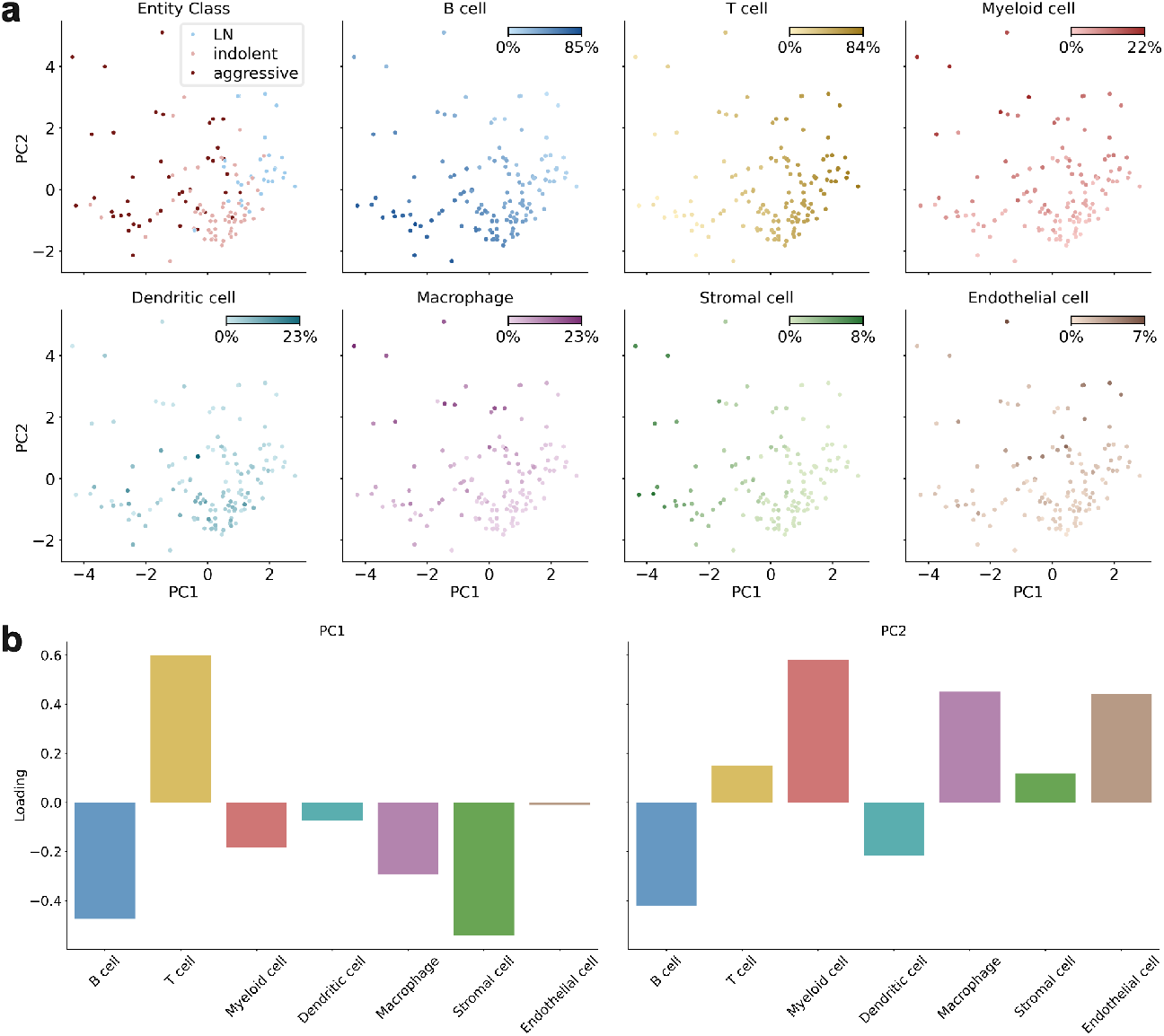
PCA of the patients in the BNHL cohort computed on the relative cell type abundances. **a**, PC1 vs PC2, colored by entity class and abundance of the major cell types. **b**, Loadings of the first two principal components show that PC1 is dominated by differences in B, T, and stromal cell composition. PC2 distinguishes based on B, myeloid, endothelial cells and macrophages.

**Supplementary Figure 4:**
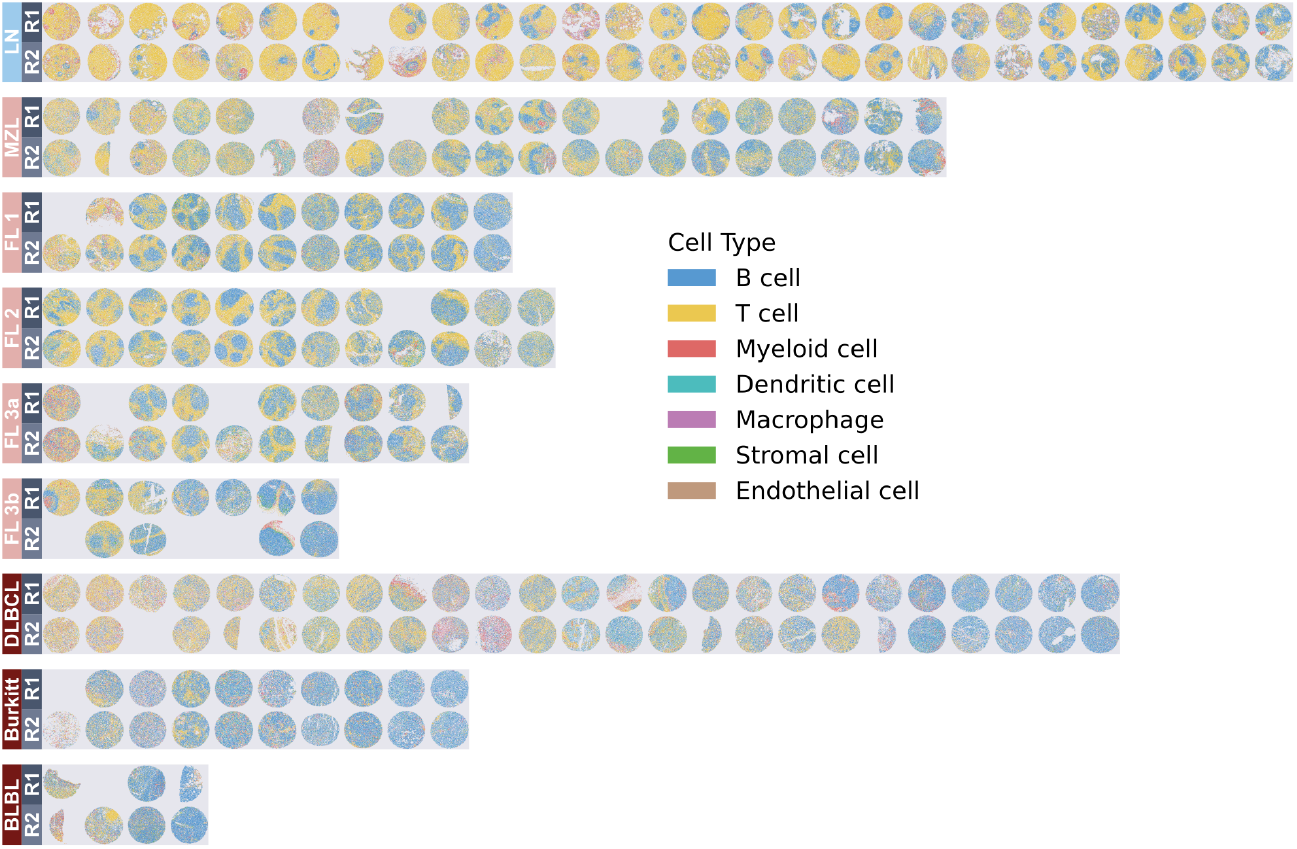
Coarse cell type predictions for all samples, stratified by entity. Replicates are shown above one another.

**Supplementary Figure 5:**
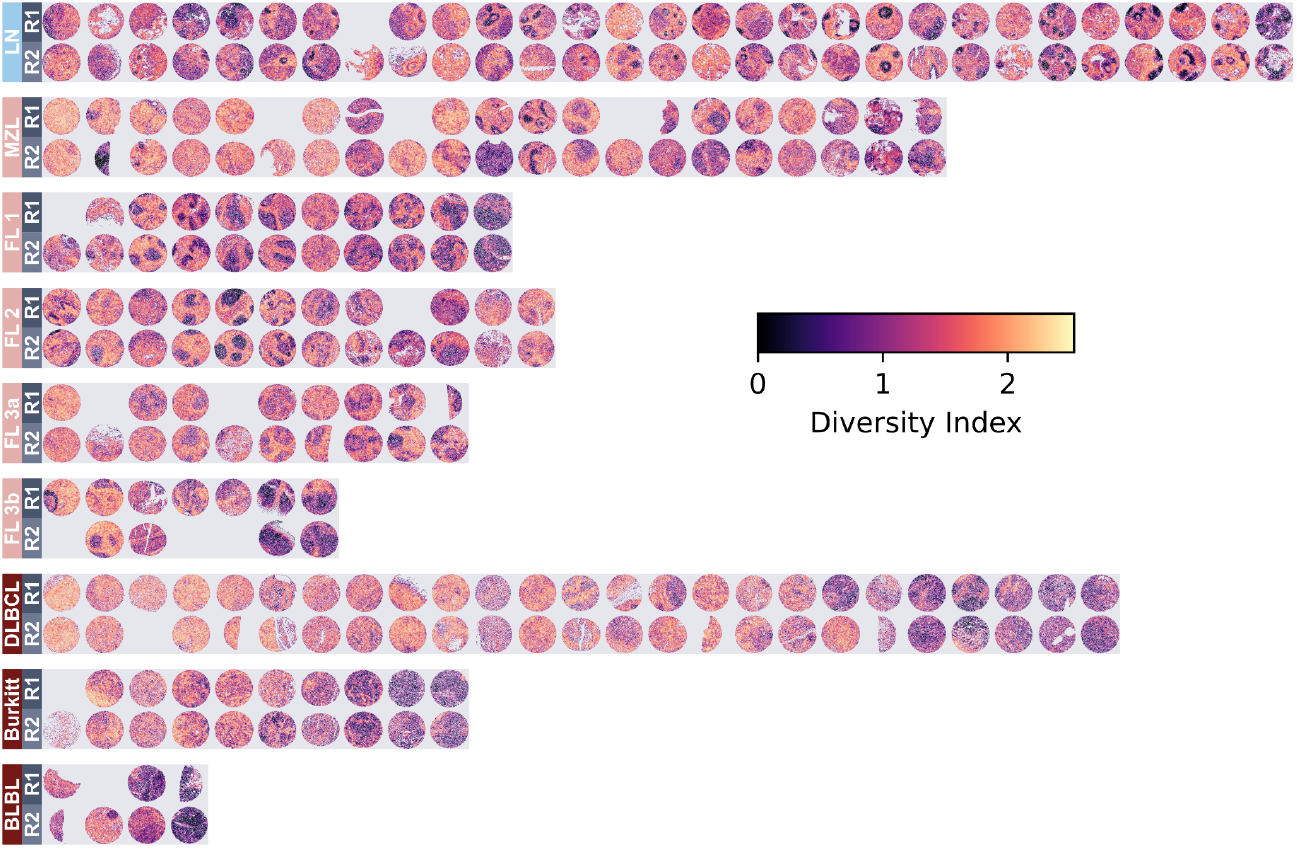
Shannon’s diversity index of a cell’s microenvironment. Due to T cells being predicted at higher granularity, T cell rich zones tend to look more heterogeneous than B cell rich zones.

**Supplementary Figure 6:**
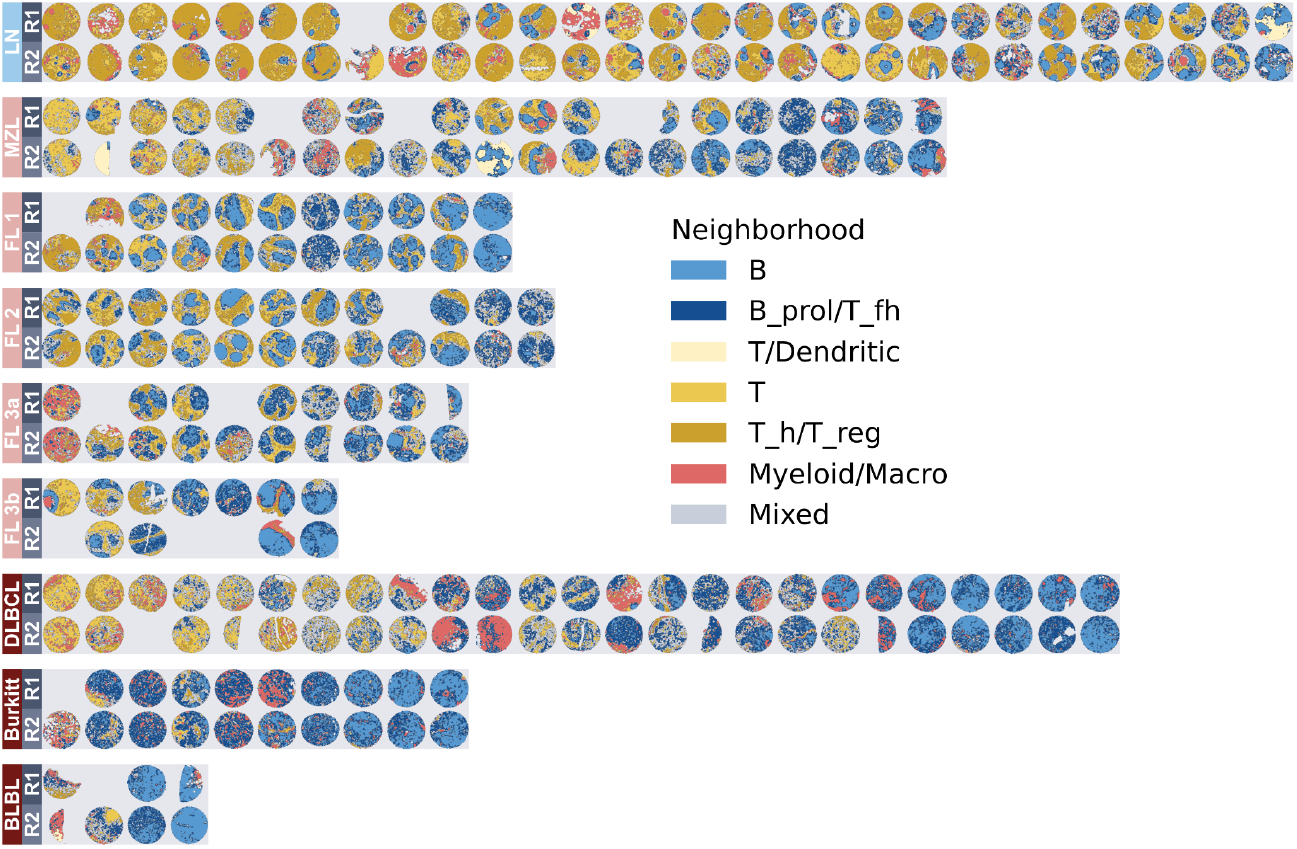
Neighborhood assignments for all samples, stratified by entity. Replicates are shown above one another.

## supplementary_tables

**Supplementary Table 1:** Overview of the TMA and WSI datasets (entity, size, number of cells per sample).

**Supplementary Table 2:** Overview of the marker panel for the TMAs.

**Supplementary Table 3:** Overview of the thresholds used for image processing and cell type classification for the TMAs.

**Supplementary Table 4:** Overview of the marker panel for the WSIs.

**Supplementary Table 5:** Overview of the thresholds used for image processing and cell type classification for the WSIs.

## References

Alaggio, R. et al. (2022) The 5th edition of the World Health Organization Classification of Haematolymphoid Tumours: Lymphoid Neoplasms. Leukemia, 36, 1720–1748.

Bird, R. (1998) Introduction to functional programming using Haskell.

Black, S. et al. (2021) CODEX multiplexed tissue imaging with DNA-conjugated antibodies. Nat. Protoc., 16, 3802–3835.

Blampey, Q. et al. (2024) Sopa: a technology-invariant pipeline for analyses of image-based spatial omics. Nat. Commun., 15, 4981.

Bortolomeazzi, M. et al. (2022) A SIMPLI (Single-cell Identification from MultiPLexed Images) approach for spatially-resolved tissue phenotyping at single-cell resolution. Nat. Commun., 13, 781.

Ding, D.Y. et al. (2025) Quantitative characterization of tissue states using multiomics and ecological spatial analysis. Nat. Genet., 57, 910–921.

Geuenich, M.J. et al. (2021) Automated assignment of cell identity from single-cell multiplexed imaging and proteomic data. Cell Syst, 12, 1173–1186.e5.

Giesen, C. et al. (2014) Highly multiplexed imaging of tumor tissues with subcellular resolution by mass cytometry. Nat. Methods, 11, 417–422.

Greenwald, N.F. et al. (2022) Whole-cell segmentation of tissue images with human-level performance using large-scale data annotation and deep learning. Nat. Biotechnol., 40, 555–565.

Hickey, J.W. et al. (2021) Strategies for accurate cell type identification in CODEX multiplexed imaging data. Front. Immunol., 12, 727626.

Hoyer, S. and Hamman, J. (2017) xarray: N-D labeled arrays and datasets in Python. Journal of open research software, 5.

Kim, J. et al. (2022) Unsupervised discovery of tissue architecture in multiplexed imaging. Nat. Methods, 19, 1653–1661.

Kinkhabwala, A. et al. (2021) MACSima imaging cyclic staining (MICS) technology reveals combinatorial target pairs for CAR T cell treatment of solid tumors. Sci. Rep., 12.

Kuswanto, W. et al. (2023) Highly multiplexed spatial profiling with CODEX: bioinformatic analysis and application in human disease. Semin. Immunopathol., 45, 145–157.

Li, N. et al. (2023) Mapping and modeling human colorectal carcinoma interactions with the tumor microenvironment. Nat. Commun., 14, 7915.

Marconato, L. et al. (2024) SpatialData: an open and universal data framework for spatial omics. Nature.

Nirmal, A.J. and Sorger, P.K. (2024) SCIMAP: A Python Toolkit for Integrated Spatial Analysis of Multiplexed Imaging Data. J Open Source Softw, 9.

Palla, G. et al. (2022) Squidpy: a scalable framework for spatial omics analysis. Nature.

Peng, T. et al. (2017) A BaSiC tool for background and shading correction of optical microscopy images. Nat. Commun., 8, 14836.

Phillips, D. et al. (2021) Highly multiplexed phenotyping of immunoregulatory proteins in the tumor microenvironment by CODEX tissue imaging. Front. Immunol., 12, 687673.

Punovuori, K. et al. (2024) Multiparameter imaging reveals clinically relevant cancer cell-stroma interaction dynamics in head and neck cancer. Cell, 187, 7267–7284.e20.

Righelli, D. et al. (2022) SpatialExperiment: infrastructure for spatially-resolved transcriptomics data in R using Bioconductor. Bioinformatics, 38, 3128–3131.

Rocklin, M. (2015) Dask: Parallel Computation with Blocked algorithms and Task Scheduling. SciPy, 126–132.

Roider, T. et al. (2024) Multimodal and spatially resolved profiling identifies distinct patterns of T cell infiltration in nodal B cell lymphoma entities. Nat. Cell Biol., 26, 478–489.

Russ, J.C. (2006) The image processing handbook, fifth edition 5th ed. CRC Press, Boca Raton, FL.

Schapiro, D. et al. (2022) MCMICRO: a scalable, modular image-processing pipeline for multiplexed tissue imaging. Nat. Methods, 19, 311–315.

Schmidt, U. et al. (2018) Cell Detection with Star-Convex Polygons. In, Medical Image Computing and Computer Assisted Intervention – MICCAI 2018. Springer International Publishing, pp. 265–273.

Schniederjohann, C. et al. (2025) Multiplexed immunophenotyping of lymphoma tissue samples. Methods Mol. Biol., 2865, 375–393.

Schürch, C.M. et al. (2020) Coordinated cellular neighborhoods orchestrate antitumoral immunity at the colorectal cancer invasive front. Cell, 182, 1341–1359.e19.

Scott, D.W. and Gascoyne, R.D. (2014) The tumour microenvironment in B cell lymphomas. Nat. Rev. Cancer, 14, 517–534.

Sofroniew, N. et al. (2024) napari: a multi-dimensional image viewer for Python Zenodo.

Stringer, C. et al. (2021) Cellpose: a generalist algorithm for cellular segmentation. Nat. Methods, 18, 100–106.

Tan, Y. et al. (2024) SPACEc: A Streamlined, Interactive Python Workflow for Multiplexed Image Processing and Analysis. bioRxiv, 2024.06.29.601349.

Varrone, M. et al. (2024) CellCharter reveals spatial cell niches associated with tissue remodeling and cell plasticity. Nat. Genet., 56, 74–84.

Verschoor, C.P. et al. (2015) An introduction to automated flow cytometry gating tools and their implementation. Front. Immunol., 6, 380.

Virshup, I. et al. (2021) anndata: Annotated data. BioRxiv.

Virshup, I. et al. (2023) The scverse project provides a computational ecosystem for single-cell omics data analysis. Nat. Biotechnol., 41, 604–606.

Windhager, J. et al. (2023) An end-to-end workflow for multiplexed image processing and analysis. Nat. Protoc., 18, 3565–3613.

